# Aggregate trait evolvability and macroevolution in two sister species of the bryozoan *Stylopoma*

**DOI:** 10.1101/2021.12.09.471970

**Authors:** Sarah Leventhal, Sarah Jamison-Todd, Carl Simpson

**Affiliations:** University of Colorado Museum of Natural History and Department of Geological Sciences, University of Colorado, Boulder 265 UCB Boulder, CO 80309; University College London, Earth Sciences, 5 Gower Place, London, WC1E 6BS

**Keywords:** Bryozoans, aggregate traits, macroevolution, quantitative genetics, trait variation

## Abstract

The study of trait evolution in modular animals is more complicated than that in solitary animals, because a single genotype of a modular colony can express an enormous range of phenotypic variation. Furthermore, traits can occur either at the module level or at the colony level. However, it is unclear how the traits at the colony level evolve. We test whether colony-level aggregate traits, defined as the summary statistics of a phenotypic distribution, can evolve. To quantify this evolutionary potential, we use parent-offspring pairs in two sister species of the bryozoan *Stylopoma*, grown and bred in a common garden breeding experiment. We find that the medians of phenotypic distributions are evolvable between generations of colonies. We also find that the structure of this evolutionary potential differs between these two species. Ancestral species align more closely with the direction of species divergence than the descendent species. This result indicates that aggregate trait evolvability can itself evolve.

## Introduction

### bryozoans as a model system in the study of aggregate trait evolution

In modular animals, such as bryozoans and siphonophores, colonies are composed of genetically identical individuals that possess a wide range of phenotypic variation (McKinney and Jackson, 1991; Lidgard, 1990; Damian-Serrano et al., 2021). This variation is expressed among colony members so that colonies represent a snapshot of the phenotypic variation that stems from a single genotype (Cheetham et al., 1995, 1993; Damian-Serrano et al., 2021). At its most extreme, this variation within colonies is expressed as morphologically discontinuous body types, called polymorphs, which serve different functions in the colony. How polymorphism in modular animals evolves is unknown (Taylor, 2020; Lidgard et al., 2012; Simpson et al., 2017), but it is impossible to resolve without first understanding whether phenotypic distributions within colonies are actually evolvable across colony generations. To understand whether phenotypic distributions are evolvable, we turn to a clade of modular animals called bryozoans.

Cheilostomes are the most diverse clade of bryozoans in the modern oceans, with over 7,000 described species (Bock, 2022). Like all bryozoans, cheilostomes have a complex life cycle with distinct asexual and sexual modes of reproduction (Taylor, 2020; McKinney and Jackson, 1991). Asexual reproduction allows for a single colony to grow larger by increasing the number of modules (zooids) in the colony. Cheilostome bryozoans sexually reproduce to form new colonies (Lidgard and Jeremy, 1989; McKinney and Jackson, 1991; Taylor, 2020). The larvae formed by sexual reproduction between colonies establish new, genetically-distinct colonies upon settling.

It is not clear whether the variation expressed within a colony is passed between generations of colonies. The evolvability of such variation would support existing explanations for the evolution of division of labor in bryozoan colonies, because polymorph types are believed to be derived from preexisting body types, termed autozooids, within colonies (McKinney and Jackson, 1991; Taylor, 2020; Treibergs and Giribet, 2020). These body types are thought to evolve gradually, so that unimodal phenotypic distributions become multimodal (Cheetham, 1973). If aspects of these distributions are evolvable, then they can presumably split and diverge over time, allowing for morphologically distinct zooids to evolve.

The phenotypic variation exhibited within a colony does not arise from microenvironmental change along axes of colony growth (Okamura, 1992). Nor is such phenotypic variation the result of positive trait correlation between asexually-budded zooids (Simpson et al., 2020). Rather, differences in phenotypes among zooids appears to be developmental in nature (Taylor, 2020; McKinney and Jackson, 1991). Yet, species of bryozoans are different from each other. For those differences to evolve, phenotypic distributions of zooid-level traits in colonies are required to be inherited across colony generations.

There are generally two types of colony-level traits: aggregate traits, which are summary statistics of traits in zooids, and emergent traits, which are expressed wholly at the colony-level (Grantham, 1995; Okasha, 2014; Lloyd and Gould, 1993). Evolutionary potential of aggregate traits has rarely been assessed, with the exception of some studies (e.g., Jablonski 1986 and Jablonski and Hunt 2006). In bryozoans, emergent colony-level traits, such as the position and orientation of polymorphs relative to other zooids in colonies, are known to have evolutionary potential (Simpson et al., 2020). While the sorting of aggregate traits has been subject to some prior study (e.g., Lloyd and Gould 1993), the evolutionary potential of aggregate traits has never been assessed. Aggregate traits are therefore the focus of our study (Fig. 1).

**Figure 1:**
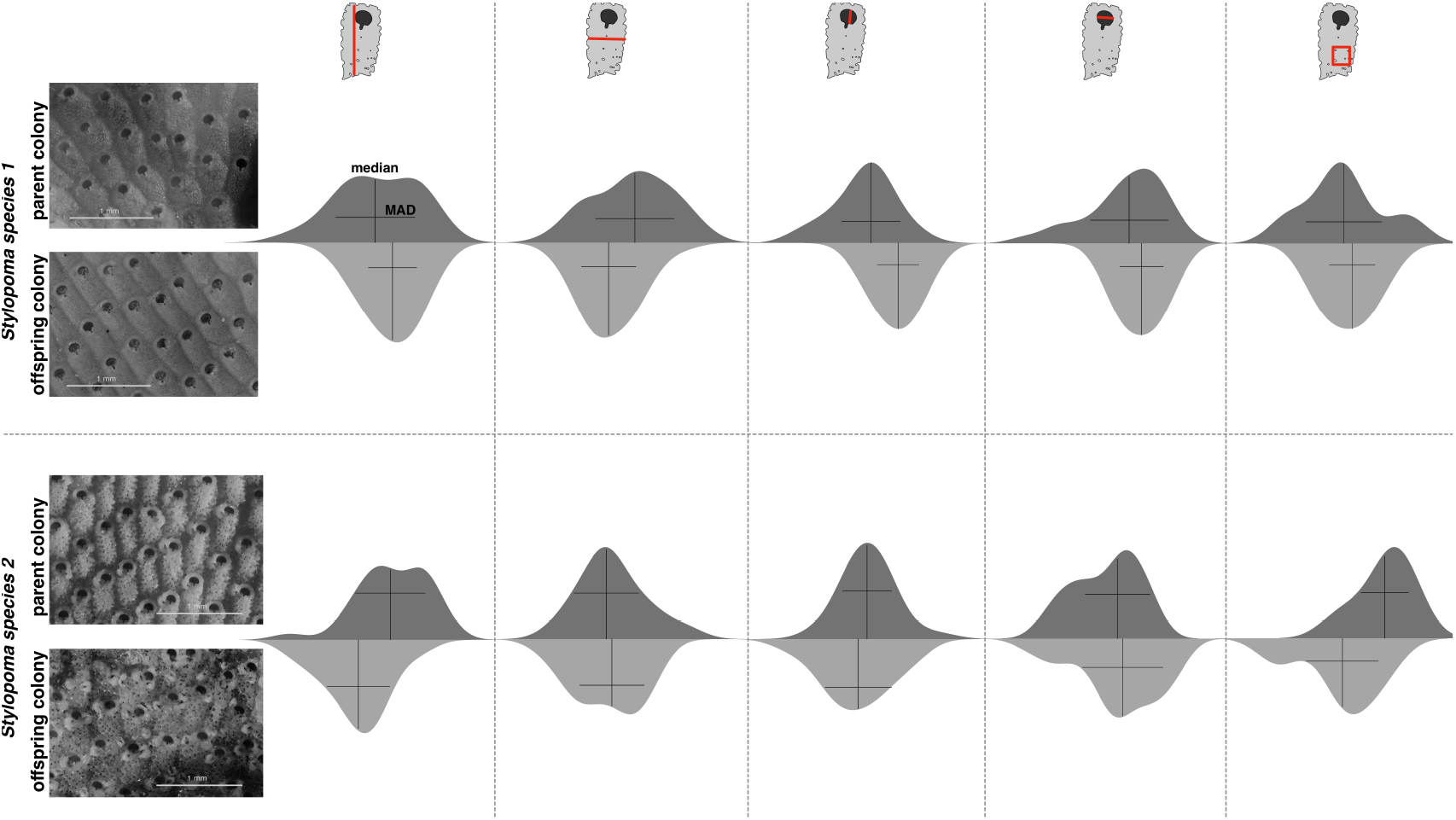
An example of aggregate phenotypic traits in parent and offspring colonies. We show examples of two colony generations from two species of *Stylopoma*. The resulting distributions of traits are displayed to the right of each maternal and offspring colony. The traits we consider are body length, body width, orifice length, orifice width, and frontal pore density, indicated by red lines. To quantify the distributions, we measure two aspects of these distributions: central tendency with the median (vertical black lines) and dispersion around the median with median absolute deviation (MAD, horizontal black lines). The analysis in this study measures parent-offspring covariation by comparing the medians and median absolute deviations of traits across colony generations.

Here we focus on two extant species of the cheilostome bryozoan *Stylopoma. Stylopoma* is a dominant member of cryptic coral reef communities (Jackson and Cheetham, 1990) with an array of polymorphs expressed in colonies (Simpson et al., 2017). The specimens we use were grown and bred in a common garden environment to evaluate whether skeletons demonstrate stability across generations (Jackson and Cheetham, 1990; Cheetham et al., 1993). Furthermore, these specimens were used to assess patterns of speciation, serving as an important model for punctuated equilibrium (Jackson and Cheetham, 1999).

### Evolvability as evolutionary potential

In this study we test the hypothesis that phenotypic distributions of zooid-level traits in colonies are evolvable. These colonies were used in a series of pioneering studies on quantitative genetics of bryozoans during the 1990s (Cheetham et al., 1993, 1994, 1995). However, these studies focused on zooid-level traits and did not consider colony-level evolution. Here we use new measurements on these specimens to quantify the evolvability of colony-level traits, and therefore their evolutionary potential.

We use evolvability, as defined by Houle (Houle, 1992) to determine whether aggregate traits in colonies can evolve. Evolvability is a measure of trait variance that is scaled by the trait mean, and can be interpreted as the maximum possible rate of evolutionary change that can occur across a single generation under direct selection (Houle, 1992; Hansen and Houle, 2008; Love et al., 2021; Jablonski, 2022).

Evolvability is difficult to quantify. It is possible to estimate it from a phenotypic variance-covariance matrix (derived from *V_P_*), which is generally considered a sufficient proxy for an underlying genetic variance-covariance matrix (Cheverud, 1988; Roff, 1995; Sodini et al., 2018; Love et al., 2021). However, while this may hold true for solitary animals, it is not applicable to colonial animals (Cheetham et al., 1993), nor is it safe to assume that *V_P_* captures the evolvability of aggregate traits. We estimate evolvability from parent-offspring covariance matrices, which capture the covariation of traits across generations (Rice, 2004). A parent-offspring covariance matrix is a better approximation of genetic variation than a simple phenotypic variance-covariance matrix, allowing for us to determine whether trait variation is underpinned by some degree of genetic variation.

Cheetham et al. (1993) found that *V_P_* and *V_G_* of zooid-level traits are not comparable and, through quantifying the *V_G_* of zooid level traits, showed that they are evolvable. We use these same specimens to estimate the evolvability of colony-level traits.

## Methods

### Specimens and image acquisition

We used colonies from two species of the bryozoan genus *Stylopoma*. These colonies were originally used in several studies on skeletal morphology and the assessment of morphological differences between species of *Stylopoma* (Jackson and Cheetham, 1990; Cheetham et al., 1993, 1994, 1995). While several unnamed species of *Stylopoma* were described in these studies, we focus on two species that were determined to be sisters: *S. species 1* and *S. species 2*. Colonies of these species were grown in the 1980s from a larval stage to adulthood in a common environment at the Smithsonian’s field station in the San Blas Islands in Panama. The experimental design was based on Maturo’s protocol (Maturo Jr, 1973) to isolate maternal colonies with a clean, flat substrate on which their larvae could settle. Once offspring colonies were established, they were reared in a common environment.

The experimental design is intended to minimize the impact of environment on phenotypic variation in colonies, so that phenotypic differences between colonies can be attributable to genetic differences. Furthermore, the flat surfaces colonies use as substrates minimize distortion of zooid morphology for measurement collection from two-dimensional images. More details on the rearing of these specimens can be found in Jackson and Cheetham 1990; Cheetham et al. 1993; Simpson et al. 2020.

We used a total of 93 parent-offspring colony pairs of *Stylopoma species 1* (42 pairs) and *Stylopoma species 2* (53 pairs).

To digitize colonies, we first used a stackshot rail, a Canon EOS camera, and Helicon Remote software to generate z-stacks of colonies. Our camera was set to 4x magnification, and we set our stackshot rail to move in increments that have 10% focus overlap between photos. We used Helicon Focus software to stitch together our image stacks, yielding high-resolution focus-stacked photographs for measurement. We evaluated this technique for error by repeatedly measuring the same zooids in a colony photographed at different angles. We found that at angles of 20° to 30°, we had a difference of ±20*μ*m in measurements of autozooid body length.

### Quantifying aggregate colony-level traits

We measured two summary statistics of the distributions of five autozooid traits in digitized parent and offspring colonies. To collect our measurements, we used ImageJ (Schneider et al., 2012). Within each colony, we selected thirty autozooids for measurement. In developing a selection schema of zooids in colonies, we had to consider colony astogeny (Mazurek, 2008). In the early stages of colony development, the phenotypic expression of zooids is substantially more variant than the range of expression found in later stages of colony growth (Taylor, 1986). This highly variant region of colonies is termed the zone of astogenetic change, and it is recognizable in colonies because of its irregular zooids (Urbanek, 2004). Once a colony reaches a certain stage of development, it transitions from the budding of highly variant zooids to much more controlled zooid-level expression, referred to as the zone of astogenetic repetition. In our sampling schema, we specifically targeted zooids in zones of astogenetic repetition so that astogenetic influence could be minimized. There is some evidence that zones of astogenetic change can recur in colonies as they age (Urbanek, 2003), so we were careful in selecting zooids that fell into highly controlled zones of astogenetic repetition, suggesting that the role of astogeny in the range of phenotypic expression that we measured is minimized.

On our thirty selected zooids, we measured four skeletal traits: zooid length, zooid width, orifice length, and orifice width. A fifth trait, frontal pore density, was quantified using a subset of five zooids per colony. We selected these traits because they are taxonomically significant in *Stylopoma*, as they allow for the morphological distinction of these two species (Simpson et al., 2020; Jackson and Cheetham, 1990). We did not consider avicularian morphology in this analysis, since the two species have different types of avicularia (vicarious in *S. species 1* and adventitious (as well as vicarious) in *S. species 2*).

To assess the measurement error for these traits, we measured each trait on a single zooid from each species ten times. The ranges of the measurements in each species were calculated and divided by the mean measured value in each species. Our error estimates were averaged between the two species, yielding single estimates of measurement error for each trait. The resulting measurement error values were 3.8% for zooid length, 4.0% for zooid width, 4.1% for orifice length, 2.8% for orifice width, and 4.9% for frontal pore density.

The colony-level traits we selected are based on two summary statistics that quantify the phenotypic distributions of constituent zooids. We first quantified the median phenotype (the value that splits the lower and higher halves of the distribution) of each trait in each colony. Then, we calculated the median absolute deviation (MAD) of each trait distribution to quantify dispersion among zooids within a colony. We chose these two summary statistics as proxies for the mean and variance because they are robust to small sample sizes (Whitley and Ball, 2001; Leys et al., 2013; Kashif et al., 2017). We confirmed the normality of our collected measurements with a Shapiro-Wilk test, which indicates that our median and MAD values are an estimate of mean and variance in each colony. Given the large number of manual measurements (> 25,000) required for this study, we prioritized measuring more colonies over sampling more zooids within a colony.

### Measuring colony-level trait evolvability

We follow Schluter (Schluter, 1996) to calculate a parent-offspring covariance matrix using parent-offspring pairs, and we follow Hansen (Hansen et al., 2003; Hansen and Houle, 2008) to calculate evolvabilities. First, we calculated the covariances of median and MAD trait values across parent and offspring colony generations to form a parent-offspring covariance matrix for each trait type (medians and MADs) (Fernandez and Miller, 1985; Rice, 2004; Falconer and Mackay, 1996). Since these colonies are sexually produced, we doubled these covariance matrices to estimate the genetic contribution of each colony-level trait (Falconer and Mackay, 1996). To deal with the natural asymmetry of a parent-offspring covariance matrix (e.g., the difference between the covariance of parent length and offspring width and that of offspring length and parent width), we took the mean of entries across the diagonal (Schluter, 1996).

We scaled entries in our parent-offspring covariance matrices by dividing them with the squared mean value for each trait (Hansen et al., 2011; Houle, 1992). For off-diagonal values, we use the product of the two trait means instead of a single trait squared mean. Scaling the parent-offspring covariance matrix in this way yields evolvabilities (Houle, 1992). Traits in our parent-offspring covariance matrices with diagonal values below zero were set to zero, since negative values indicate that there is no genetic variation in those traits (Cheetham et al., 1995).

For each species, we generated 2 sets of evolvability matrices: one set considers MAD and median traits separately, and the other considers MAD and median traits pooled together. We estimate matrices of median and MAD traits separately for the purpose of comparing evolvability matrices across trait type, which is not possible using a concatenated matrix. However, we also want to quantify the covariation between trait types, which requires a matrix that considers the covariation of median traits with MAD traits.

We calculated the standard error in our estimates of trait evolvability using the method outlined by Lynch and Marsh (Lynch and Walsh, 1998). To follow this method, we computed the variance of the ratio of trait variances and means (Lynch and Walsh, 1998; Garcia-Gonzalez et al., 2012). We first calculated the large sample variance for each trait mean and variance, and then computed the variance of their ratio. We took the square root of the resulting variance and divided it by the square root of sample size to estimate the standard error of each evolvability estimate.

### Evolvability comparisons and repeatability estimation

We compare our evolvabilities in two different ways: between species and across trait type. We use two different statistical tests to evaluate structural similarity between matrices for our comparisons: random skewers (Cheverud, 1996; Cheverud and Marroig, 2007) and Krzanowski correlations (Krzanowski, 1993). All statistical tests were performed in RStudio 2021.09.0 (Team, 2015) using the EvolQG package (Melo et al., 2015).

The random skewers method estimates the degree of similarity between two covariance matrices by finding the correlation of their responses to multiplication with random vectors (Cheverud, 1996). These random vectors are generated from a uniform distribution of values, which are normalized to a length of one to simulate a random selection gradient (Marroig and Cheverud, 2007). Using this method, we generated 10,000 random vectors and used the mean correlation of the resulting response vectors from each matrix as an estimate of matrix similarity.

Krzanowski correlations measure shared space by summing square vector correlations between the first *k* principal components of each matrix, where 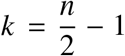 for any *n* × *n* matrix (Krzanowski, 1993; Melo et al., 2015). The resulting sum of correlations ranges from 0 to 1, with 0 indicating no similarity and 1 indicating perfect similarity. This test has no associated significance test, so we use it as a tool to assess the fractional similarity between matrices.

To estimate sampling error in our matrices, we calculate the repeatability of each matrix using a bootstrap procedure (Marroig and Cheverud, 2007; de Oliveira et al., 2009). We sampled values from our dataset with replacement 1,000 times, and used these resampled values to compute covariance matrices. The mean correlation of our observed covariance matrices with these bootstrap-generated matrices is the repeatability of each covariance matrix (Melo et al., 2015). We then used the random skewers method to compare our resampled datasets to our original dataset. The mean correlations represent the repeatability of our matrices.

### Quantifying directions of evolutionary change

To calculate trait divergence between species, we first pooled the species together and calculated the pooled mean for each trait. Then we divided the mean trait values for each species by the pooled mean for each trait. We subtracted the scaled median and MAD trait means of *Stylopoma species 2* from those of *Stylopoma species 1* to estimate divergence for each trait (Hopkins, 2016). We compute two divergence vectors: one for median traits and one for MAD traits.

### Estimating trait-specific patterns of species divergence

Since these two species of *Stylopoma* are sisters (Jackson and Cheetham, 1994), we have the opportunity to understand how closely the structure of genetic variation in each species aligns with each trait dimension. In our case, we want to evaluate the angle of deviation between the first two eigenvectors of each evolvability matrix and the dimensional axis of each trait. For convenience, we refer to these first two eigenvectors as 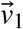 and 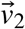, respectively. 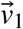 corresponds to the major axis of variation in each evolvability matrix, while 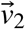 corresponds to the second major axis of variation in each matrix. These two eigenvectors capture 99.6% of the variation in median values and 94.7% of the variation in MAD values in *S. species 1*, and 99.9% of the variation in median values and 99.8% of the variation in MAD values in *S. species 2*. We therefore estimate the angle between the 2 first eigenvectors of each matrix (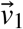 and 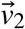) and a vector of each trait dimension.

For this part of the analysis, we use an expanded trait dimension vector, with an equal number of rows and columns, where each diagonal entry corresponds to a different trait dimension. To estimate angles, we computed the arccosine of the dot product of 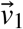 and 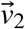 and each row of the corresponding dimension vector. The resulting angles represent the deviation between 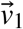 and 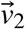 and the dimensional axis of each trait (Fig. 2). A small angle of deviation with a trait dimension indicates that there is high phenotypic variance in that trait dimension, while a large angle of deviation indicates that there is low phenotypic variance in that trait dimension.

**Figure 2:**
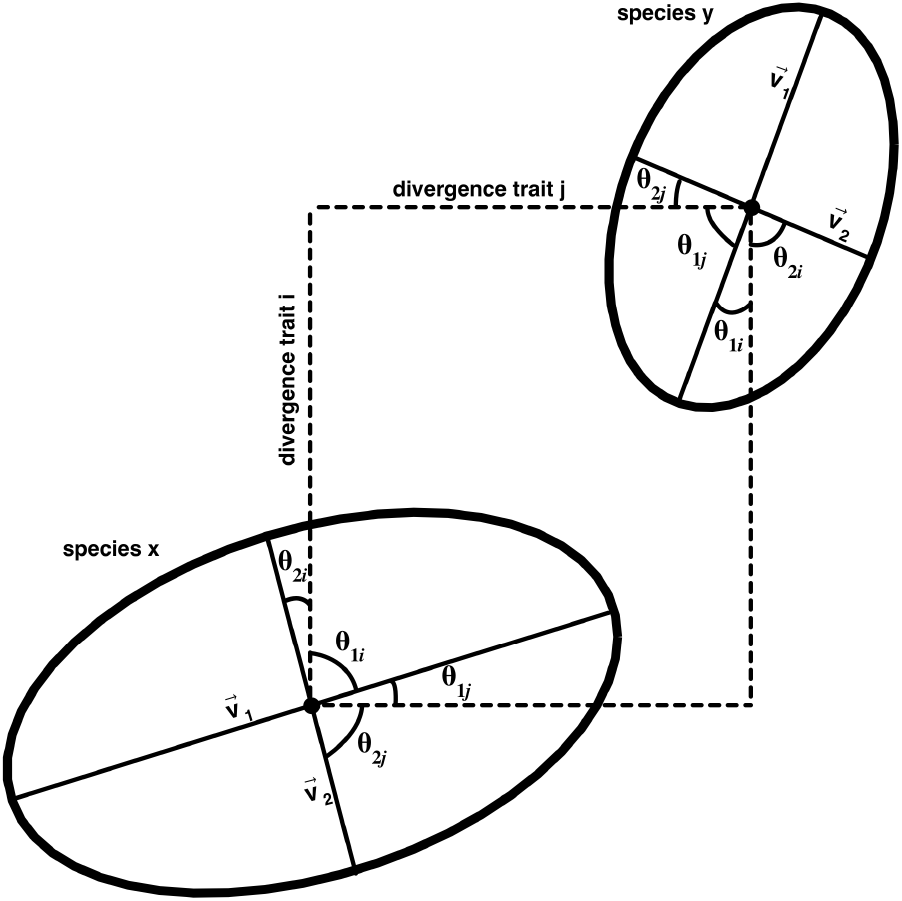
Schematic figure of our deviation angle analysis. Each ellipse corresponds to the evolvability matrix of species x and species y, simplified in two dimensions, *i* and *j*. We estimate the angles between the first two eigenvectos of each ellipse, 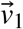 and 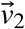, respectively, and each trait dimension, *i* and *j*. In the scenario of evolution along genetic lines of least resistance, we would expect more divergent traits to be more closely aligned with 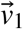 and 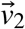 in each species.

The resulting angles provide insight into how closely 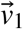 and 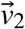 in each species aligns with each trait dimension. We compared the angles (which we call *θ_i_*, for traits 1 through *i*) calculated from the procedure above with the corresponding trait divergence between species to see if traits more closely aligned with eigenvectors in each species are also more divergent between species. For each comparison, we calculated Pearson’s r coefficients to assess the significance of the relationship.

### Estimating null values

While our repeatability estimates range from 0.95 to 0.97 for all of our calculated matrices, repeatability is a metric sensitive to sample size, with small sizes yielding inflated repeatability (Cheverud and Marroig, 2007). To estimate null evolvability values for comparison with our estimated values, we used a randomization procedure (Roff et al., 2012). We generated 1,000 randomized parent-offspring covariance matrices with replacement, and doubled these matrices and averaged them across the diagonals so that they would be symmetrical. To estimate null evolvabilities, we divided the diagonal values by the mean trait values squared, following the same methodology as we did for estimating evolvabilities from our measured parent-offspring covariance matrices (Schluter, 1996). We also calculated the standard deviation of our randomized evolvability estimates, and used them to construct 95% confidence intervals.

We assessed the significance of our calculated angles of deviation between trait-specific species divergence and the first eigenvector of our evolvability matrices by generating a distribution of angles computed between 10,000 randomly-generated vector pairs. We computed the mean and standard deviation of this distribution to compare with our calculated angles.

## Results and discussion

### The partial evolvability of colony-level phenotypic distributions

We find that median trait values of phenotypic distributions are largely evolvable in both species, with more nuanced results for MAD traits (Tables 1 and 2). Notably, the median trait values of orifice width were found to have no genetic covariation for either species. For our MAD evolvabilities, each species has different traits with evolvabilities of zero (stemming from the corresponding parent-offspring covariance values falling below zero): for *Stylopoma species 1*, the MADs of pore density and orifice length have genetic variations of zero, and for *Stylopoma species 2*, the MADs of body length and body width have genetic variations of zero.

**Table 1:** Trait evolvabilities, with error and null estimates, for *Stylopoma species 1*. Evolvability values for five trait medians and five trait MADs. ol=orifice length, ow=orifice width, pd=pore density, evol. = evolvability.

| | evolvability | standard error | Mean Null evol. ±95% confidence intervals |
| --- | --- | --- | --- |
| median length | 0.0019 | $\pm 0.00026$ | $-0.000036 \pm 0.000079$ |
| median width | 0.0053 | $\pm 0.00031$ | $-0.000095 \pm 0.00026$ |
| median ol | 0.0023 | $\pm 0.00043$ | $0.0000063 \pm 0.00018$ |
| median ow | 0.00 | $\pm 0.00040$ | $0.0000068 \pm 0.00014$ |
| median pd | 0.017 | $\pm 0.0013$ | $0.00016 \pm 0.0014$ |
| MAD length | 0.0065 | $\pm 0.0047$ | $0.000066 \pm 0.0019$ |
| MAD width | 0.018 | $\pm 0.0042$ | $-0.00017 \pm 0.0027$ |
| MAD ol | 0.00 | $\pm 0.0048$ | $0.00021 \pm 0.0018$ |
| MAD ow | 0.010 | $\pm 0.0057$ | $0.00057 \pm 0.0017$ |
| MAD pd | 0.00 | $\pm 0.030$ | $-0.0058 \pm 0.017$ |

**Table 2:** Trait evolvabilities, with error and null estimates, for *Stylopoma species 2*. Evolvability values for five trait medians and five trait MADs. ol=orifice length, ow=orifice width, pd=pore density, evol. = evolvability.

| | parent-offspring covariance evol. | standard sample error | Null evol. $\pm 95\%$ confidence intervals |
| --- | --- | --- | --- |
| median length | 0.0021 | $\pm 0.00039$ | $-0.000055 \pm 0.00023$ |
| median width | 0.0013 | $\pm 0.00041$ | $-0.000025 \pm 0.00015$ |
| median ol | 0.0019 | $\pm 0.00032$ | $0.0000095 \pm 0.000059$ |
| median ow | 0.00 | $\pm 0.00020$ | $0.000023 \pm 0.000034$ |
| median pd | 0.073 | $\pm 0.00041$ | $-0.00055 \pm 0.0019$ |
| MAD length | 0.00 | $\pm 0.0055$ | $-0.00067 \pm 0.0015$ |
| MAD width | 0.00 | $\pm 0.023$ | $-0.00063 \pm 0.0019$ |
| MAD ol | 0.021 | $\pm 0.0060$ | $-0.00015 \pm 0.00083$ |
| MAD ow | 0.044 | $\pm 0.012$ | $-0.00069 \pm 0.0030$ |
| MAD pd | 0.16 | $\pm 0.033$ | $-0.0012 \pm 0.0045$ |

The mean evolvability of nonzero MAD values is higher than that of the medians in both species (0.0090 versus 0.0021, respectively, in *S. species 1*; 0.027 and 0.0048, respectively, in *S. species 2*), but MAD evolvabilities also have larger error, likely due to the similarity in MAD values between unrelated colonies. It is possible that the controlled environment in which colonies were reared may have impacted the range of phenotypic variation expressed, thereby increasing the phenotypic similarity expressed among colonies within each species, and artificially reducing our estimated evolvabilities for MAD traits. Prior studies have found that estimates of genetic and phenotypic variation can change depending on environmental plasticity (Bégin and Roff, 2001). In our case, general phenotypic similarities across all colonies may be the result of similar environmental influence (Bull, 1987).

Regardless of its source, the large error in MAD traits indicates that some nonzero MAD evolvabilities may not be significantly above the null evolvability value. In contrast, all nonzero median evolvabilities are significantly above the null evolvability value. Furthermore, there are more MAD traits with no evolvability than there are median traits (Tables 1 and 2). This indicates that evolution of the central location of a colony-level phenotypic distribution in morphospace may evolve more readily than its dispersion around the median. In other words, aggregate traits within colonies primarily evolve according to translational shifts of the median.

### Connecting aggregate traits to division of labor

The evolution of polymorphism involves the diverging and splitting of phenotypic distributions over evolutionary timescales, which can presumably occur as a random process or a result of selective pressure (Banta et al., 1973; Silen, 1977). Translational shifts in medians of trait distributions, which our results strongly support, may underpin part of this evolutionary process. However, full understanding of this evolutionary transition requires a deeper understanding of how the shape of phenotypic distributions (i.e., trait MADs) can evolve. We explore some of these possibilities below.

There are some MAD traits that have evolvability values significantly above the null value: body width and orifice width in *S. species 1* and pore density and orifice width in *S. species 2*. The non-null evolvabilities of these traits suggests that they can potentially respond to direct selection. While the evolvability of trait variation is not a novel concept (Bruijning et al., 2020; Hill and Mulder, 2010), it has never before been studied with respect to aggregate traits. Our results suggest that phenotypic variation can itself be an evolvable trait in colonial animals, thereby indicating that the shape of a phenotypic distribution, and not just its location in morphospace, is able to respond to selection.

The non-null evolvability of both median and MAD values for some of our measured phenotypic distributions, like that of width in *S. species 1* and that of pore density in *S. species 2*, is significant because it indicates that the first two moments of a phenotypic distribution can both respond to selection. Furthermore, our concatenated covariance matrices reveal that median and MAD traits have some degree of covariation (Tables 3-4). Selection on some median traits may lead to changes in variation of other traits, and vice-versa. MAD traits that have nonzero but nonsignificant evolvability values can potentially still evolve through their covariance with more evolvable traits. For example, MAD of orifice width in *S. species 2* is close to the null evolvability value when considering standard error, which indicates it may not have significant evolvability. However, the MAD of orifice width has covariation with traits that are significantly evolvable, like median pore density. Thus, traits with null evolvabilities may still be able to evolve through their covariation with other highly evolvable traits.

**Table 3:** Evolvability matrix for *S. species 1*. med=median, mad= MAD, ol=orifice length, ow=orifice width, pd=pore density.

|  | med length | med width | med ol | med ow | med pd | mad length | mad width | mad ol | mad ow | mad pd |
| --- | --- | --- | --- | --- | --- | --- | --- | --- | --- | --- |
| med length | 0.00194 | 0.00263 | 0.00075 | 0.00000 | -0.00041 | 0.00424 | 0.00123 | 0.00000 | -0.00278 | 0.00000 |
| med width |  | 0.00528 | 0.00187 | 0.00000 | -0.00343 | 0.00144 | -0.00026 | 0.00000 | -0.00605 | 0.00000 |
| med ol |  |  | 0.00225 | 0.00000 | 0.00220 | 0.00993 | 0.01088 | 0.00000 | -0.00605 | 0.00000 |
| med ow |  |  |  | 0.00000 | 0.00000 | 0.00000 | 0.00000 | 0.00000 | 0.00000 | 0.00000 |
| med pd |  |  |  |  | 0.01720 | -0.00985 | 0.00300 | 0.00000 | -0.00100 | 0.00000 |
| mad length |  |  |  |  |  | 0.00654 | 0.01566 | 0.00000 | -0.00033 | 0.00000 |
| mad width |  |  |  |  |  |  | 0.01811 | 0.00000 | 0.00779 | 0.00000 |
| mad ol |  |  |  |  |  |  |  | 0.00000 | 0.00000 | 0.00000 |
| mad ow |  |  |  |  |  |  |  |  | 0.01018 | 0.00000 |
| mad pd |  |  |  |  |  |  |  |  |  | 0.00000 |

**Table 4:** Evolvability matrix for *S. species 2*. med=median, mad= MAD, ol=orifice length, ow=orifice width, pd=pore density.

|  | med length | med width | med ol | med ow | med pd | mad length | mad width | mad ol | mad ow | mad pd |
| --- | --- | --- | --- | --- | --- | --- | --- | --- | --- | --- |
| med length | 0.00213 | 0.00065 | -0.00006 | 0.00000 | -0.00222 | 0.00000 | 0.00000 | -0.00640 | 0.00560 | -0.00833 |
| med width |  | 0.00133 | -0.00036 | 0.00000 | -0.00225 | 0.00000 | 0.00000 | 0.00103 | -0.00619 | -0.00313 |
| med ol |  |  | 0.00190 | 0.00000 | 0.00399 | 0.00000 | 0.00000 | 0.00458 | 0.00545 | -0.01098 |
| med ow |  |  |  | 0.00000 | 0.00000 | 0.00000 | 0.00000 | 0.00000 | 0.00000 | 0.00000 |
| med pd |  |  |  |  | 0.07253 | 0.00000 | 0.00000 | 0.03264 | 0.02457 | 0.09083 |
| mad length |  |  |  |  |  | 0.00000 | 0.00000 | 0.00000 | 0.00000 | 0.00000 |
| mad width |  |  |  |  |  |  | 0.00000 | 0.00000 | 0.00000 | 0.00000 |
| mad ol I |  |  |  |  |  |  |  | 0.02076 | 0.03823 | -0.00889 |
| mad ow |  |  |  |  |  |  |  |  | 0.04414 | -0.02023 |
| mad pd |  |  |  |  |  |  |  |  |  | 0.15696 |

### Trait divergence and the evolution of evolvability

The comparison of our evolvability matrices between species varies depending on the trait type. For median traits, the evolvability matrices for *S. species 1* and *S. species 2* are similar (Table 1, Random skewers Pearson’s *r* = 0.801, *p* < 0.05; Krzanowski Pearson’s *r* = 0.756). However, for MAD trait evolvabilities, the matrices are unrelated (Random skewers Pearson’s *r* = 0.106, *p* ~ 0.4; Krzanowski Pearson’s *r* = 0.290). This indicates that the evolvabilities of MAD traits in these two species are likely significantly different. Thus, the major axes of variation for MAD traits appear to have evolved since *S. species 1* and *S. species 2* diverged (Arnold et al., 2008).

If evolvabilities have been conserved in each species since *S. species 1* and *S. species 2* diverged, we would expect 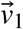 and 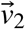, which are the first two eigenvectors of each evolvability matrix, to have similar angles of deviation from each trait dimension. While this holds true for median trait evolvabilities (Fig. 3 A, B), this is not the case for MAD trait evolvabilities (Fig 2 C, D). This result is unsurprising, since we find evolvability matrices for MAD values are divergent between species. However, to understand how this divergence between MAD evolvability matrices emerged, we can compare correlations of trait-specific species divergence and the angles of deviation between the first two eigenvectors of each evolvability matrix and each trait dimension.

**Figure 3:**
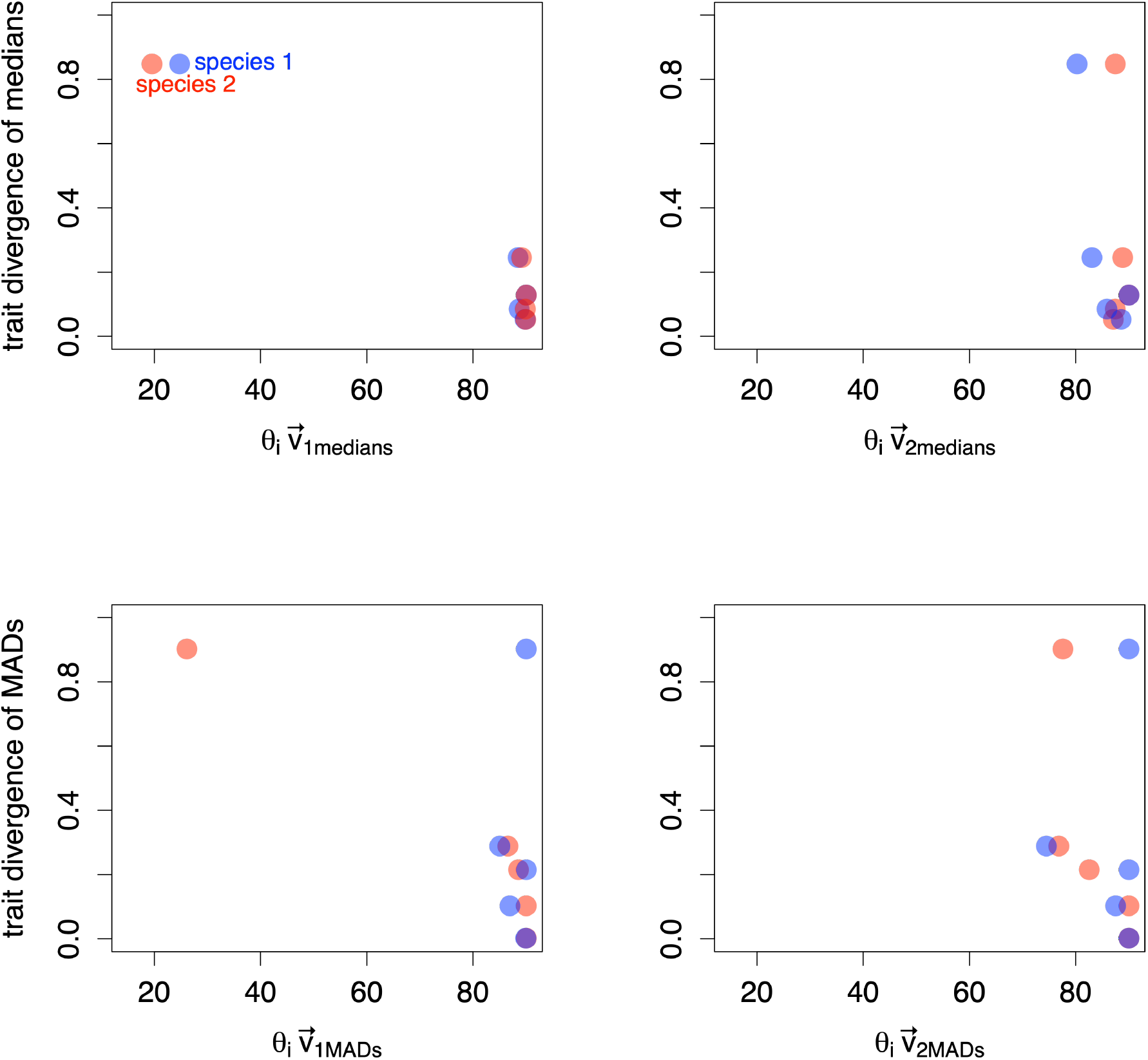
Deviations between axes of variation and trait dimensions plotted against corresponding trait divergence. **A:** Angle between each median trait dimension and 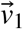 of median trait evolvabilities plotted against the trait divergence vector (Pearson’s *r* = −0.978 for *S. species 1;* Pearson’s *r* = −0.977 for *S. species 2*). **B:** Angle between each median trait dimension and 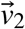 of median trait evolvabilities in each species plotted against the trait divergence vector (Pearson’s *r* = −0.827 for *S. species 1* and Pearson’s *r* = −0.19 for *S. species 2*). **C:** Angle between each MAD trait dimension and 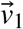 of MAD trait evolvabilities plotted against the trait divergence vector (Pearson’s *r* = 0.21 for *S. species 1*, Pearson’s *r* = −0.964 for *S. species 2*. **D:** Angle between each MAD trait dimension and 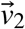 of MAD trait evolvabilities plotted against the trait divergence vector Pearson’s *r* = 0.075 for *S. species 1* and Pearson’s *r* = −0.731 for *S. species 2*

We find that the angles of deviation between 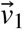 and 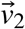 of MAD traits and each MAD trait dimension in *S. species 2* have a negative correlation with corresponding trait divergences between species (Pearson’s *r* = −0.964 for 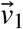 and Pearson’s *r* = −0.731 for 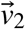). Additionally, in both species, we find that there is a negative correlation between trait divergence of medians and the angle of deviation between the first two eigenvectors of trait median evolvability and each trait dimension (Pearson’s *r* = −0.978 for 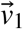 and Pearson’s *r* = −0.827 for 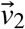 for *S. species 1*; Pearson’s *r* = −0.977 for 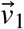 and Pearson’s *r* = −0.19 for 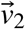 in *S. species 2*). In other words, we find that trait dimensions which are closely aligned to 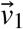 and 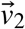 are also more divergent between species. This finding connects our microevolutionary observation of trait covariation with the macroevolutionary pattern of trait divergence between species.

It is worth noting that trait dimensions are, in many cases, nearly orthogonal to 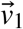 and 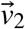. We anticipated the correlations for 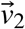 to be weaker than those for 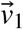, but even in our estimates of 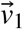, most angles are close to 90° (Fig. 3 A-D). Our computed negative correlations are largely driven by a single character that is highly divergent between species: frontal pore density. For both the median and MAD trait evolvabilities of *S. species 2* and the median evolvabilities of *S. species 1*, frontal pore density is the trait dimension most closely aligned with 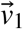 and 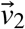. Furthermore, it is the trait with the highest component loading for the first principal component of *S. species 1* median trait evolvabilities, and of *S. species 2* median and MAD trait evolvabilities (Table 4). Thus, it is not surprising that the median and MAD values of this trait comprise much of the structure of variation.

We expected to see a negative correlation between the trait divergence and the deviation of 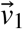 from each trait dimension, since that indicates that most evolution occurs in traits with the largest contributions to 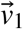. This represents a trait-specific version of evolution along genetic lines of least resistance, which has been readily observed in a wide array of solitary animals (Schluter, 1996; Hunt, 2007; Hopkins et al., 2016). Therefore, it appears that the structure of variation in median traits for *S. species 1* and median and MAD traits in *S. species 2* follows the expected pattern of trait divergence along evolutionary lines of least resistance.

However, 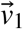 and 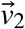 of the MAD trait evolvability matrix in *S. species 1* do not align most closely with the most divergent traits between species (Fig. 3 C, D). Instead, 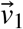 and 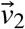 of MAD evolvabilities in *S. species 1* appear to have a slightly positive correlation with pattern of trait divergence, as the angles between each eigenvector and a vector of trait dimensions show no correlation with trait divergence (Pearson’s *r* = 0.21 for 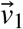 and Pearson’s *r* = 0.075 for 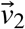). This indicates that less divergent MAD traits have more variation in *S. species 1*. Furthermore, the angles of deviation between each eigenvector and each trait dimension fall close to our mean null angle value of 74.9°, which indicates that there is likely no relationship between MAD trait evolvability in *S. species 1* and the pattern of divergence between species. This pattern is largely driven by the lack of genetic variation for the MAD of frontal pore density in this species, which is highly divergent between species (Table 6). However, there is no genetic covariation of this trait across colony generations, making it non-evolvable in this species.

**Table 5:** Matrix Comparisons. Here we show the results of our matrix comparisons between species and traits. Pearsons’s *r* values are shown for each statistical test. When applicable, significant similarity values are indicated by *. evol. = evolvability matrix, sp1 = *S. species 1*, sp2 = *S. species 2*, med = median, MAD = median absolute deviation

| Matrix 1 | Matrix 2 | Random skewers | Krzanowski Correlation |
| --- | --- | --- | --- |
| evol. sp1 med | evol. sp2 med | 0.801* | 0.756 |
| evol. sp1 MAD | evol. sp2 MAD | 0.106 | 0.290 |
| evol. sp1 med | evol. sp1 MAD | 0.301 | 0.407 |
| evol. sp2 med | evol. sp2 MAD | 0.761* | 0.500 |

**Table 6:** First two principal components of parent-offspring evolvability matrices. Here we show the eigenvalues and component loadings corresponding to each trait for the first two principal components of our four evolvability matrices. sp 1 med = evolvability matrix for median traits of *S. species 1*, sp 1 mad = evolvability matrix for MAD traits of *S. species 1*, sp 2 med = evolvability matrix for median traits of *S. species 2*, sp 2 mad = evolvability matrix for MAD traits of *S. species 2*, ol= orifice length, ow = orifice width, pd = pore density.

|  | sp 1 med |  | sp1 mad |  | sp 2 med |  | sp 2 mad |  |
| --- | --- | --- | --- | --- | --- | --- | --- | --- |
|  | PC1 | PC2 | PC1 | PC2 | PC1 | PC2 | PC1 | PC2 |
| Eigenvalue | 0.018 | 0.0071 | 0.031 | 0.0098 | 0.073 | 0.0024 | 0.16 | 0.067 |
| length | -0.060 | -0.44 | -0.51 | -0.42 | -0.032 | 0.88 | 0 | 0 |
| width | -0.25 | -0.76 | -0.81 | -0.067 | -0.032 | 0.47 | 0 | 0 |
| ol | 0.099 | -0.44 | 0 | 0 | 0.056 | -0.075 | -0.12 | 0.59 |
| ow | 0 | 0 | -0.29 | 0.91 | 0 | 0 | -0.20 | 0.77 |
| pd | 0.96 | -0.179 | 0 | 0 | 0.99 | 0.047 | 0.97 | 0.23 |

*1* median trait evolvabilities, and of *S. species 2* median and MAD trait evolvabilities (Table 4).

Fossils show that *S. species 1* diverged from *S. species 2* approximately 3 million years ago (Jackson and Cheetham, 1999). Our results indicate that speciation occurred according to the variance structure of the ancestral species (*S. species 2*), with that of the descendent species (*S. species 1)* (Jackson and Cheetham, 1999) possibly diverging after speciation. Instability of genetic variations on macroevolutionary timescales has been observed previously (Roff, 2000; Arnold et al., 2008). Whether aggregate traits in colonies are less stable than traits in solitary animals on geologic timescales is still unclear. At the very least, the case of MAD traits in *Stylopoma* appears to be an example in which aggregate trait evolvabilities themselves evolve (Pigliucci, 2008). The evolving evolvability of aggregate traits may explain the relatively rapid trait evolution observed in some bryozoan taxa (Di Martino and Liow, 2022). If trait evolvabilities change frequently, then new axes of morphological variation can emerge as colonies evolve. On macroevolutionary timescales, this can lead to unexpected patterns of species divergence and, potentially, the emergence of novel morphological adaptations and features (Wagner and Draghi, 2010).

## Conclusions

In this study, we have investigated aggregate trait evolvability in two sister species of the bryozoan *Stylopoma*. There are several key takeaways that we want to highlight. We find that aggregate traits evolve in bryozoans. Our results suggest that phenotypic distributions evolve through translational shifts in morphospace. Still, some phenotypic distributions can evolve through changes to trait dispersion, providing an avenue for the evolution of trait variation. MAD trait evolvabilities also appear more unstable than those of median traits, indicating that they may change more readily on macroevolutionary timescales. This brings us to our final point, which is that the structure of variation in colonies can evolve, and that changes to this variation can contribute to evolution of aggregate trait distributions.

How changes to evolvability, especially in aggregate traits, may contribute to directions of speciation is a topic worthy of future investigation, since bryozoans appear to undergo morphological evolution on geologically rapid timescales. The key to understanding how modular animals like bryozoans express and maintain such a wide array of morphologies in colonies is the study of how their aggregate traits and underlying evolvabilities evolve and may underpin the evolution of polymorphism.

The evolution of polymorphism remains one of the greatest unknowns in the study of colonial animals. The evolutionary potential of aggregate traits provides one avenue for polymorphism to evolve, as the development of polymorphs stems from the splitting of phenotypic distributions of traits in colonies. Since such phenotypic distributions are evolvable, the study of how they shift and change shape on macroevolutionary timescales is the key to understanding the emergence of division of labor in colonies.

## Supporting information

https://datadryad.org/stash/share/X_plYI0csh6d1yvdbOp8H4b9HiqN-R2apPuTn_nCCQw

## Acknowledgements

We thank J.B.C. Jackson and A. Herrera-Cubilla for collecting and breeding the specimens used in this experiment. We thank J. Li and A. Halling for providing manuscript revision. This research was supported by the Penny Patterson Graduate Fellowship Fund and the Bruce and Marcy Benson Graduate Student Fellowship. The original breeding experiments conducted in 1989 to 1990 were supported by the Smithsonian Institution Scholarly Studies Program and by the Smithsonian Tropical Research Institute.

## Author Contributions

S.L. and S.J-T. digitized all available specimens for analysis and collected measurement data.

S.L. performed the data analysis. S.L. and C.S. wrote the manuscript.

## Competing Interests Statement

The authors declare that they have no competing interests.

## Data Accessibility

All data needed to evaluate the conclusions in the paper are present in the paper and/or Supplementary Materials. Additional data related to this paper may be requested from the authors.

## Notes

### Competing Interest Statement

The authors have declared no competing interest.

### Summary of Updates

Figure 1 has been revised and a new figure has been added (now Figure 2). The text has been revised to introduce bryozoans, aggregate traits, and evolvability more clearly.

## References and Notes

Arnold, S. J., Bürger, R., Hohenlohe, P. A., Ajie, B. C., and Jones, A. G. (2008). Understanding the evolution and stability of the g-matrix. Evolution, 62(10):2451–2461.

Banta, W. C., Boardman, R., Cheetham, A., and Oliver, W. (1973). Evolution of avicularia in cheilostome Bryozoa, pages 295–303. Van Nostrand Reinhold, 1 edition.

Bock, P. (2022). Stylopoma levinsen, 1909.

Bruijning, M., Metcalf, C. J. E., Jongejans, E., and Ayroles, J. F. (2020). The evolution of variance control. Trends in Ecology Evolution, 35(1):22–33.

Bull, J. J. (1987). Evolution of phenotypic variance. Evolution, 41(2):303–315.

Bégin, M. and Roff, D. A. (2001). An analysis of g matrix variation in two closely related cricket species, gryllus firmus and g. pennsylvanicus. Journal of Evolutionary Biology, 14(1):1–13.

Cheetham, A. H. (1973). Study of cheilostome polymorphism using principal components analysis. Living and Fossil Bryozoa. Academic Press, London, pages 385–409.

Cheetham, A. H., Jackson, J. B. C., and Hayek, L.-A. C. (1994). Quantitative genetics of bryozoan phenotypic evolution. ii. analysis of selection and random change in fossil species using reconstructed genetic parameters. Evolution, 48(2):360–375.

Cheetham, A. H., Jackson, J. B. C., and Hayek, L.-A. C. (1995). Quantitative genetics of bryozoan phenotypic evolution. iii. phenotypic plasticity and the maintenance of genetic variation. Evolution, 49(2):290–296.

Cheetham, A. H., Jackson, J. B. C., and Hayek, L. C. (1993). Quantitative genetics of bryozoan phenotypic evolution. i. rate tests for random change versus selection in differentiation of living species. Evolution, 47(5):1526–1538.

Cheverud, J. M. (1988). A comparison of genetic and phenotypic correlations. Evolution, 42(5):958–968.

Cheverud, J. M. (1996). Quantitative genetic analysis of cranial morphology in the cotton-top (saguinus oedipus) and saddle-back (s. fuscicollis) tamarins. Journal of Evolutionary Biology, 9(1):5–42.

Cheverud, J. M. and Marroig, G. (2007). Research article comparing covariance matrices: random skewers method compared to the common principal components model. Genetics and Molecular Biology, 30:461–469.

Damian-Serrano, A., Haddock, S. H. D., and Dunn, C. W. (2021). The evolution of siphonophore tentilla for specialized prey capture in the open ocean. Proceedings of the National Academy of Sciences, 118(8):e2005063118.

de Oliveira, F. B., Porto, A., and Marroig, G. (2009). Covariance structure in the skull of catarrhini: a case of pattern stasis and magnitude evolution. Journal of Human Evolution, 56(4):417–430.

Di Martino, E. and Liow, L. H. (2022). Changing allometric relationships among fossil and recent populations in two colonial species. Evolution.

Falconer, D. and Mackay, T. (1996). Introduction to quantitative genetics, volume 3. Harlow, Essex, UK: Longmans Green.

Fernandez, G. and Miller, J. (1985). Estimation of heritability by parent-offspring regression. Theoretical and Applied Genetics, 70(6):650–654.

Garcia-Gonzalez, F., Simmons, L. W., Tomkins, J. L., Kotiaho, J. S., and Evans, J. P. (2012). Comparing evolvabilities: common errors surrounding the calculation and use of coefficients of additive genetic variation. Evolution: International Journal of Organic Evolution, 66(8):2341–2349.

Grantham, T. A. (1995). Hierarchical approaches to macroevolution: Recent work on species selection and the “effect hypothesis”. Annual Review of Ecology and Systematics, 26(1):301–321.

Hansen, T. F. and Houle, D. (2008). Measuring and comparing evolvability and constraint in multivariate characters. Journal of Evolutionary Biology, 21(5):1201–1219.

Hansen, T. F., Pélabon, C., Armbruster, W. S., and Carlson, M. L. (2003). Evolvability and genetic constraint in dalechampia blossoms: components of variance and measures of evolvability. J Evol Biol, 16(4):754–66.

Hansen, T. F., Pélabon, C., and Houle, D. (2011). Heritability is not evolvability. Evolutionary Biology, 38(3):258.

Hill, W. G. and Mulder, H. A. (2010). Genetic analysis of environmental variation. Genetics Research, 92(5-6):381–395.

Hopkins, M. J. (2016). Magnitude versus direction of change and the contribution of macroevolutionary trends to morphological disparity. Biological Journal of the Linnean Society, 118(1):116–130.

Hopkins, M. J., Haber, A., and Thurman, C. L. (2016). Constraints on geographic variation in fiddler crabs (ocypodidae: Uca) from the western atlantic. Journal of Evolutionary Biology, 29(8):1553–1568.

Houle, D. (1992). Comparing evolvability and variability of quantitative traits. Genetics, 130(1):195–204.

Hunt, G. (2007). Evolutionary divergence in directions of high phenotypic variance in the ostracode genus Poseidonamicus. Evolution, 61(7):1560–1576.

Jablonski, D. (1986). Larval ecology and macroevolution in marine invertebrates. Bulletin of marine science, 39(2):565–587.

Jablonski, D. (2022). Evolvability and macroevolution: Overview and synthesis. Evolutionary Biology, 49(3):265–291.

Jablonski, D. and Hunt, G. (2006). Larval ecology, geographic range, and species survivorship in cretaceous mollusks: organismic versus species-level explanations. The American Naturalist, 168(4):556–564.

Jackson, J. B. C. and Cheetham, A. H. (1990). Evolutionary significance of morphospecies: A test with cheilostome bryozoa. Science, 248(4955):579.

Jackson, J. B. C. and Cheetham, A. H. (1994). Phylogeny reconstruction and the tempo of speciation in cheilostome bryozoa. Paleobiology, 20(4):407–423.

Jackson, J. B. C. and Cheetham, A. H. (1999). Tempo and mode of speciation in the sea. Trends in Ecology and Evolution, 14(2):72–77.

Kashif, M., Aslam, M., Al-Marshadi, A. H., Jun, C., and Khan, M. I. (2017). Evaluation of modified non-normal process capability index and its bootstrap confidence intervals. IEEE Access, 5:12135–12142.

Krzanowski, W. J. (1993). Permutational tests for correlation matrices. Statistics and Computing, 3(1):37–44.

Leys, C., Ley, C., Klein, O., Bernard, P., and Licata, L. (2013). Detecting outliers: Do not use standard deviation around the mean, use absolute deviation around the median. Journal of Experimental Social Psychology, 49(4):764–766.

Lidgard, S. (1990). Growth in encrusting cheilostome bryozoans: Ii. circum-atlantic distribution patterns. Paleobiology, 16(3):304–321.

Lidgard, S., Carter, M. C., Dick, M. H., Gordon, D. P., and Ostrovsky, A. N. (2012). Division of labor and recurrent evolution of polymorphisms in a group of colonial animals. Evolutionary Ecology, 26(2):233–257.

Lidgard, S. and Jeremy, B. C. J. (1989). Growth in encrusting cheilostome bryozoans: I. evolutionary trends. Paleobiology, 15(3):255–282.

Lloyd, E. A. and Gould, S. J. (1993). Species selection on variability. Proceedings of the National Academy of Sciences of the United States of America, 90(2):595–599.

Love, A. C., Grabowski, M., Houle, D., Liow, L. H., Porto, A., Tsuboi, M., Voje, K. L., and Hunt, G. (2021). Evolvability in the fossil record. Paleobiology, pages 1–24.

Lynch, M. and Walsh, B. (1998). Genetics and analysis of quantitative traits. Sinauer Associates, Inc., Sunderland, MA.

Marroig, G. and Cheverud, J. M. (2007). A comparison of phenotypic variation and covariation patterns and the role of phylogeny, ecology, and ontogeny during cranial evolution of new world monkeys. Evolution, 55(12):2576–2600.

Maturo Jr, F. (1973). Offspring variation from known maternal stocks of parasmittina nitida (verrill). Living and Fossil Bryozoa, GP Larwood, Eds.(Academic Press, 1973), pages 577–584.

Mazurek, D. (2008). A model explaining some bryozoan colonies. Nature Precedings.

McKinney, F. and Jackson, J. (1991). Bryozoan Evolution. University of Chicago Press.

Melo, D., Garcia, G., Hubbe, A., Assis, A. P., and Marroig, G. (2015). Evolqg - an r package for evolutionary quantitative genetics [version 3; referees: 2 approved, 1 approved with reservations]. F1000Research, 4(925).

Okamura, B. (1992). Microhabitat variation and patterns of colony growth and feeding in a marine bryozoan. Ecology, 73(4):1502–1513.

Okasha, S. (2014). Emergent group traits, reproduction, and levels of selection. Behavioral and Brain Sciences, 37(3):268–269.

Pigliucci, M. (2008). Is evolvability evolvable? Nature Reviews Genetics, 9(1):75–82.

Rice, S. (2004). Evolutionary Theory: Mathematical and Conceptual Foundations. Sinauer.

Roff, D. (2000). The evolution of the g matrix: selection or drift? Heredity, 84(2):135–142.

Roff, D. A. (1995). The estimation of genetic correlations from phenotypic correlations: a test of cheverud’s conjecture. Heredity, 74:481–490.

Roff, D. A., Prokkola, J. M., Krams, I., and Rantala, M. J. (2012). There is more than one way to skin a g matrix. Journal of Evolutionary Biology, 25(6):1113–1126.

Schluter, D. (1996). Adaptive radiation along genetic lines of least resistance. Evolution, 50(5):1766–1774.

Schneider, C. A., Rasband, W. S., and Eliceiri, K. W. (2012). NIH Image to ImageJ: 25 years of image analysis.

Silen, L. (1977). Polymorphism, book section 6, pages 183–231. Academic Press.

Simpson, C., Herrera-Cubilla, A., and Jackson, J. B. C. (2020). How colonial animals evolve. Science Advances, 6(2).

Simpson, C., Jackson, J. B. C., and Herrera-Cubilla, A. (2017). Evolutionary determinants of morphological polymorphism in colonial animals. The American Naturalist, 190(1):17–28.

Sodini, S. M., Kemper, K. E., Wray, N. R., and Trzaskowski, M. (2018). Comparison of genotypic and phenotypic correlations: Cheverud’s conjecture in humans. Genetics, 209(3):941–948.

Taylor, P. (2020). Bryozoan Paleobiology. Wiley.

Taylor, P. D. (1986). The ancestrula and early growth pattern in two primitive cheilostome bryozoans: Pyripora catenularia (fleming) and pyriporopsis portlandensis pohowsky. Journal of Natural History, 20(1):101–110.

Team, R. (2015). Rstudio: integrated development for r. rstudio. Inc., Boston, MA, 700.

Treibergs, K. A. and Giribet, G. (2020). Differential gene expression between polymorphic zooids of the marine bryozoan bugulina stolonifera. G3 (Bethesda), 10(10):3843–3857.

Urbanek, A. (2003). Organization and evolution of animal colonies. Biology Bulletin of the Russian Academy of Sciences, 30(1):1–8.

Urbanek, A. (2004). Morphogenetic gradients in graptolites and bryozoans. Acta Palaeontologica Polonica, 49(4).

Wagner, G. P. and Draghi, J. (2010). Evolution of Evolvability.

Whitley, E. and Ball, J. (2001). Statistics review 1: Presenting and summarising data. Critical Care, 6(1):66.

